# Auditory hemispheric asymmetry as a specialization for actions and objects

**DOI:** 10.1101/2023.04.19.537361

**Authors:** Paul Robert, Robert Zatorre, Akanksha Gupta, Julien Sein, Jean-Luc Anton, Pascal Belin, Etienne Thoret, Benjamin Morillon

## Abstract

What is the function of auditory hemispheric asymmetry? We propose that the identification of sound sources relies on two complementary and perceptually relevant acoustic invariants — actions and objects — that are processed asymmetrically and sufficient to model and categorize any sound. We observed that environmental sounds are an independent combination of purely temporal and spectral acoustic modulations. Behaviorally, discrimination of actions relied on temporal modulations, while discrimination of objects relied on spectral modulations. Functional magnetic resonance imaging data showed that actions and objects are respectively decoded in the left and right hemispheres, in bilateral superior temporal and left inferior frontal regions. This asymmetry reflects a generic differential processing — through differential neural sensitivity to temporal and spectral modulations present in all environmental sounds — that supports the efficient categorization of actions and objects. These results provide an ecologically valid framework of the functional role of auditory brain asymmetry.

## Introduction

Sounds are in essence temporal signals that are continuously mapped on the basis of their spectral content. This implicit time-frequency process offers a challenge to the auditory system. A mathematical constraint, the Gabor-Heisenberg uncertainty principle, indeed implies that there is a trade-off between the precision that can be achieved in the time and frequency dimensions when analyzing an acoustic event (Gabor, 1946; Joos, 1948; Zatorre et al., 2002). However, the human perceptual system has been argued to beat this uncertainty principle (Ronken, 1971; Moore, 1973).

It has been proposed that auditory hemispheric asymmetry is an evolutionary solution that optimizes the processing of sound sources. With two hemispheres, two complementary strategies can theoretically be used to analyze sounds in parallel - respectively optimizing the temporal or spectral resolution (Zatorre et al., 2002; Washington & Tillinghast, 2015) - which would reduce redundancy and increase the efficiency of neural coding (Barlow 1961; Smith & Lewicki 2006; Gervain & Geffen, 2019). To evaluate the evidence supporting this solution at each of David Marr’s levels of analysis - implementational, algorithmic, and functional - we examine the physical realization, computational processes, and overall purpose or goal of the auditory information-processing system (Marr, 1982).

Empirical evidence shows that in humans the two auditory cortices are structurally different (Heschl, 1878; Geschwind & Levitsky, 1968; Wada, 1975). The left auditory cortex is overall bigger than its right counterpart (Penhune et al., 1996; Dorsaint-Pierre et al., 2006; Anderson et al., 1999; Meyer et al., 2014; Dalboni da Rocha et al., 2020), a trait already present in neonates (Williams et al., 2023) and shared between Old World monkeys, apes, and humans (Marie et al., 2018; Becker & Meguerditchian, 2022). It also has larger layer III magno-pyramidal cells (Hayes & Lewis, 1996; Hutsler, 2003; Hutsler & Gazzaniga, 1996), higher density of dendrites, axons and synaptic contacts (Ocklenburg et al., 2018), a larger width and number of cortical micro-columns that are also more widely spaced apart (Seldon, 1981; Chance et al., 2006), less inter-connected macro-columns (Galuske et al., 2000), more developed myelination (Anderson et al., 1999), and different network configuration and embedding within the connectome (Cha et al., 2016; Mišíc et al., 2018).

All of this suggests a faster action potential transmission in the left auditory cortex, leading to a greater temporal resolution. This is confirmed in mice with reports of shorter temporal integration windows in left auditory superficial layers, a functional asymmetry associated with weaker recurrent excitatory synaptic connectivity (Neophytou et al., 2022) that results in different circuit-motifs (Levy et al., 2019).

Accordingly, in humans, varying acoustic stimuli independently in temporal or spectral dimensions differentially drive auditory responses in the left and right hemispheres, respectively. This phenomenon is observed whether the stimuli are pure-tone sequences alternating in pitch at different temporal rates (Zatorre & Belin, 2001; Jamison et al., 2006), systematically varying noise bands in spectral width and temporal rate of change (Schönwiesner et al., 2005), or concatenated narrowband noise stimuli varying in segment transition rates (Boemio et al., 2005). The asymmetric pattern observed is consistently lateralized to the left for temporal variations and to the right for spectral variations.

These results are accounted for by the “spectrotemporal modulation (STM) asymmetry hypothesis”. It argues that the left hemisphere is capable of integrating a wide range of temporal modulations, ranging from slow to fast, but has a limited ability to integrate high spectral modulations. In contrast, its right homolog can integrate a wide range of spectral modulations, ranging from low to high, but only slow temporal modulations (Flinker et al., 2019; Figure 1A). In other words, the left and right auditory systems would respectively be sensitive to *purely* temporal (i.e., low spectral / all temporal) and *purely* spectral (i.e., slow temporal / all spectral) modulations, and not be differentially sensitive to their interaction, i.e. combined fast temporal and high spectral modulations. This formulation provides a computationally specific stimulus space (modulation domain) linked to an implementational computation in the cortex (integration of neuronal inputs from subpopulations).

**Figure 1:**
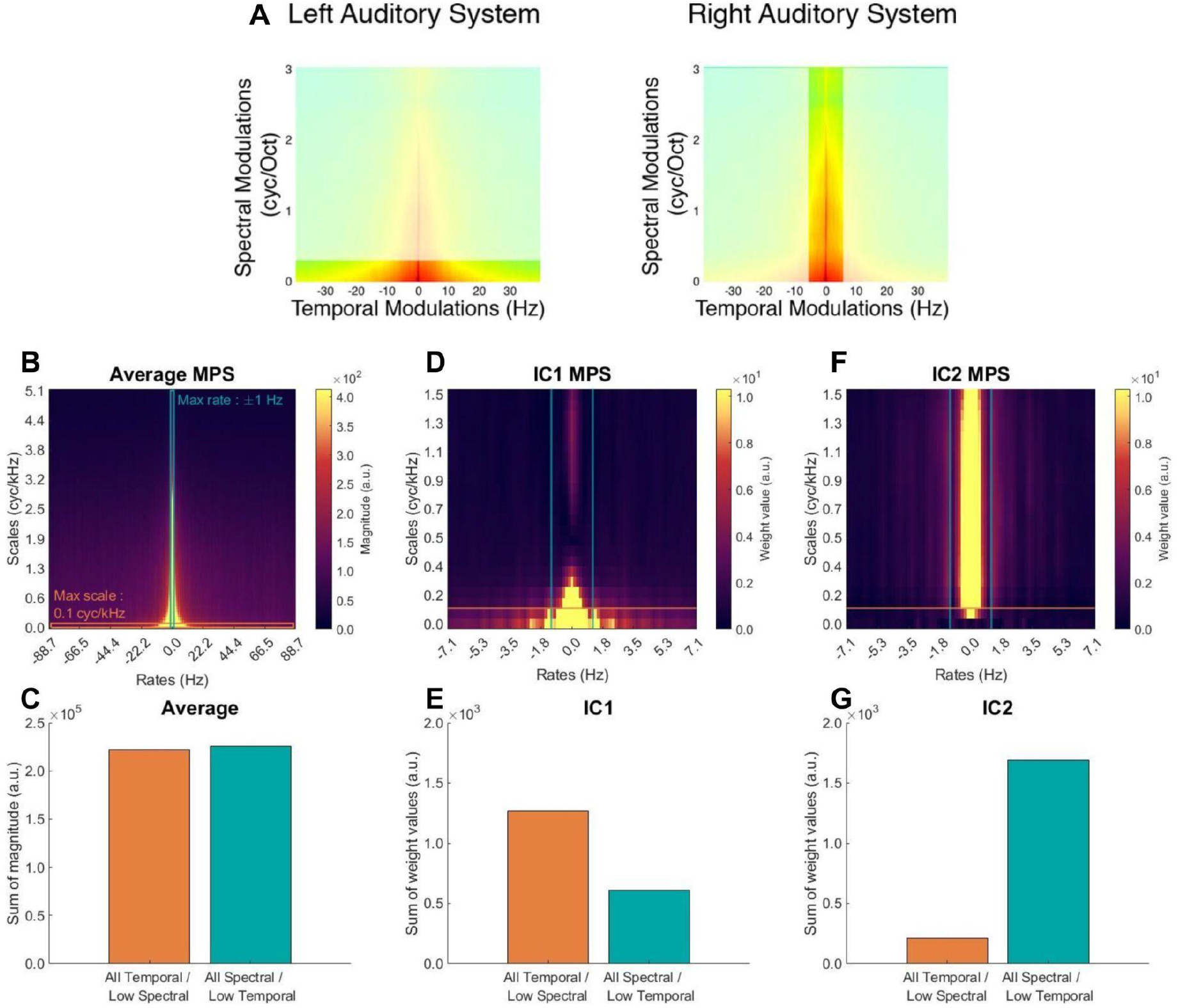
Acoustic analysis of environmental sounds. **(A)** Modulation asymmetry hypothesis, from Flinker and colleagues (2019). The left hemisphere is sensitive to purely temporal modulations while the right hemisphere is sensitive to purely spectral modulations. **(B)** Average modulation power spectrum (MPS) of environmental sounds (McDermott & Simoncelli 2011; Norman-Haignere et al., 2015). The orange subspace covers purely temporal modulations (i.e., all temporal, low spectral, < 0.1 cyc/kHz), while the blue subspace covers purely spectral modulations (i.e., all spectral, slow temporal, < 1 Hz). MPS parameters were defined so that the two subspaces have an equal summed magnitude (see B). **(C)** Summed magnitude per subspace. **(D)** First and **(F)** second independent components (IC) of the IC analysis computed across all environmental sound MPSs. Scales and rates axes are zoomed for visual clarity purposes (same conventions as in A). **(E & G)** Summed weights per subspace, for **(E)** IC1 and **(G)** IC2.

This framework is grounded in electrophysiological evidence in animal models, showing that cells in the subcortical auditory pathway and in the primary auditory cortex respond selectively to a narrow range of temporal modulations (sound amplitude modulations over time, expressed in Hz) and spectral modulations (modulations along the frequency axis in a spectrogram, expressed in cycle/kHz) (Fritz et al., 2003; Chi et al., 2005; Rodríguez et al., 2010). In humans, similar maps of spectrotemporal sensitivity are observed in the auditory cortices (Schönwiesner & Zatorre, 2009; Santoro et al., 2014, Norman-Haignere et al, 2015).

This framework was also applied to explain speech and melody perception (Albouy et al., 2020). The authors generated a corpus of *a-capella* songs for which verbal and melodic content was crossed and balanced, allowing the dissociation of speech/melody-specific cues from non-specific acoustic features. Each stimulus was then selectively filtered in either the temporal or spectral modulation dimension. Behavioral results (across both French- and English-speaking listeners) indicated that speech perception depends more on temporal cues (Shannon et al., 1995), while melody recognition relies more on spectral cues. Functional MRI (fMRI) data showed that the neural decoding of speech and melodies depends on activity patterns in the left and right associative auditory regions, respectively. Finally, degradation of the acoustic signal affected neural responses in the same way as behavioral responses. The triple dissociation between cognitive domain (speech vs melody), degradation type (temporal vs spectral), and hemisphere (left vs right) observed in this study thus strongly supports the conclusion that distinct hemispheric sensitivity to STM underlies the lateralized responses to speech and melodies.

While the STM account of hemispheric asymmetries provides a cogent and powerful framework, there has been limited integration between this model and an understanding of the characteristics of sounds from the environment for which it is presumed to be specialized. To fully appreciate this relationship, it is important to consider how sounds are naturally produced in the environment. The acoustic structure of sounds can be modeled as the result of a source being filtered by an object, such as someone knocking on a wooden door. This source-filter approach underlies the analysis and synthesis of speech (Klatt & Klatt, 1990), music (Smith, 2004), and continuous-interaction sounds (Thoret et al., 2014; Conan et al., 2014b). Additionally, it forms the foundation for understanding sound source recognition in the ecological approach to perception (Gibson, 2014; Michaels & Carello, 1981; Gaver, 1993b), where invariant structures in the signal are used for the categorization of sound sources.

The action-object framework (Conan et al., 2014b) is a formalization of the source-filter approach at the signal-processing level. It models all sounds as the output of an excitation —the action— applied to a filter —the object. The auditory system processes two categories of invariants to identify the sound source (Gaver, 1993a): transformational, which characterize the properties of the action such as its dynamics (Thoret et al., 2014; Conan et al., 2014b); and structural, which characterize the properties of the object such as its shape, size, surface properties or material (Aramaki et al., 2009; Aramaki et al., 2010; Giordano & McAdams, 2006). This distinction would appear to align with the STM framework, but direct evidence is lacking.

The transformational invariants refer to the temporal characteristics of sound such as the spectral centroid variations of friction sounds (Thoret et al., 2014) or the statistics of impacts (Conan et al., 2014b), both of which contribute to temporal modulations. The structural invariants instead refer to the spectral properties of sound such as the modal distribution of the frequencies of a vibrating object (Aramaki et al., 2011), or its surface properties when rubbed or rolled on (Conan et al., 2014a), and relate to spectral modulations. In the case of speech or musical instruments, the object represents the spectral properties of the sound, primarily driven by the vibration of the vocal folds or the resonator (oral cavity, body of instrument, etc), while the action represents the temporal evolution of this spectral content, mainly referring to prosody and linguistic rhythms in speech or the excitation in the case of musical instruments (e.g. bow on a string, vibrating reed on a column of air, etc). In sum, action and object depend on the acoustic cues that are processed asymmetrically by the auditory system.

Moreover, transformational and structural acoustic invariants are both necessary and sufficient for recognizing different action and object properties (Conan et al., 2014a; Aramaki et al., 2011; Grassi, 2005; Lemaitre & Heller, 2013; Lemaitre et al., 2018) or reenacting human actions, e.g., footsteps (Young et al., 2013), or drawing movements (Thoret et al., 2014). This ecological framework is relevant for characterizing sound signal properties that arise from physical phenomena, enabling efficient auditory recognition. Consequently, the action-object framework directly supports the functional relevance of a processing system dedicated to independently process these two acoustic invariants, making it a plausible explanation of auditory hemispheric asymmetry at the functional level.

In the present study, we sought to demonstrate that the acoustical cues pertaining to actions and objects are processed asymmetrically, which is necessary to account for auditory hemispheric asymmetry at all levels of Marrian analysis (Marr, 1982). The hemispheric dominance observed for speech and music processing could be reinterpreted as a result of each hemisphere’s specialization for two categories of invariant acoustic cues - actions and objects - found in all sounds, making it an ecologically valid explanation that aligns with the theory of efficient neural coding (Barlow 1961, Smith & Lewicki 2006; Gervain & Geffen, 2019).

To approach this goal, we first studied the acoustic structure of corpora of natural environmental sounds to estimate the validity of the STM asymmetry hypothesis. We investigated whether natural sounds can be described as an independent combination of purely temporal and purely spectral modulations. We next synthesized environmental sounds under the action/object framework to manipulate independently their temporal and spectral content. We created sounds that would result from friction motions (*actions*) on different surfaces (*objects)*. Importantly, many environmental sounds are not periodic and do not induce a pitch percept. We hence generated non-periodic sounds. Third, in a behavioral experiment we tested whether action perception relies on purely temporal modulations, while object perception relies on purely spectral modulations by degrading spectral vs temporal acoustical cues. Finally, in an fMRI experiment, we investigated whether the neural decoding of actions and objects depends on activity patterns in the left and right hemispheres, respectively.

## Results

### 1. Environmental sounds are composed of an independent combination of purely temporal and purely spectral modulations

We first investigated the acoustic structure of a corpus of natural environmental sounds, composed of a large set (N = 250) of frequently heard and recognizable sounds that humans regularly encounter (McDermott & Simoncelli, 2011; Norman-Haignere et al., 2015, Figure 1B-C). An independent component analysis (Hyvarinen, 1999) in the spectrotemporal modulation (STM) domain showed that environmental sounds can be decomposed into two independent components, explaining the majority of variance (r² = 0.801). The first component corresponds to *purely* temporal modulations (all temporal, low spectral, < 0.1 cyc/kHz; Figure 1D-E), while the second component covers *purely* spectral modulations (all spectral, slow temporal, < 1 Hz; Figure 1F-G). Most of the information in environmental sounds is hence carried by these two independent patterns of STMs, and crucially, not by a combination of fast temporal and high spectral modulations, supporting the STM framework.

### 2. Perception of actions and objects depends on purely temporal and purely spectral modulations respectively

We then synthesized environmental sounds to allow us to manipulate the temporal and spectral dimensions of each stimulus independently, an operation that would not be possible with real-life recordings. We used the *action-object* synthesis paradigm (Conan et al., 2014a) to create sounds that would result from friction motions (*actions*) on different surfaces (*objects;* Suppl. Figure S1). These stimuli have previously been used in experiments where subjects had to recognize geometric shapes (circle, oval, or line) from friction sounds produced by drawing movements (Thoret et al., 2014), and are interpreted here as a subset of real-life environmental sounds carrying cognitive (action and object domains) information.

We first created nine stimuli, by generating three distinct actions (drawing dynamics with natural motion trajectory in the 1-5 Hz range) and three distinct objects (impulse responses with surface roughness properties leading to spectral modulations below 5 cycles/kHz) and combining them together (Suppl. Figure S2). In short, the synthesis method produces friction sounds by emulating a series of impacts on a surface, with the impact density being controlled by the velocity of the movement and each impact reflecting the spectral profile of the object (Figure 2A; see Method). We computed the cepstrum amplitude of our stimuli to estimate their degree of periodicity and confirmed that they only contained residual periodicity, less than sounds known to induce a pitch percept and a right-lateralized auditory response (Suppl. Figure S3; Albouy et al., 2020). To alter these sounds in the modulation domain, we created two additional versions of each stimulus using the same thresholds as in the corpus analysis: the purely temporal, in which all spectral modulations above 0.1 cycle/kHz are filtered out, and the purely spectral, in which all temporal modulations above 1Hz are filtered out (Figure 2B).

**Figure 2:**
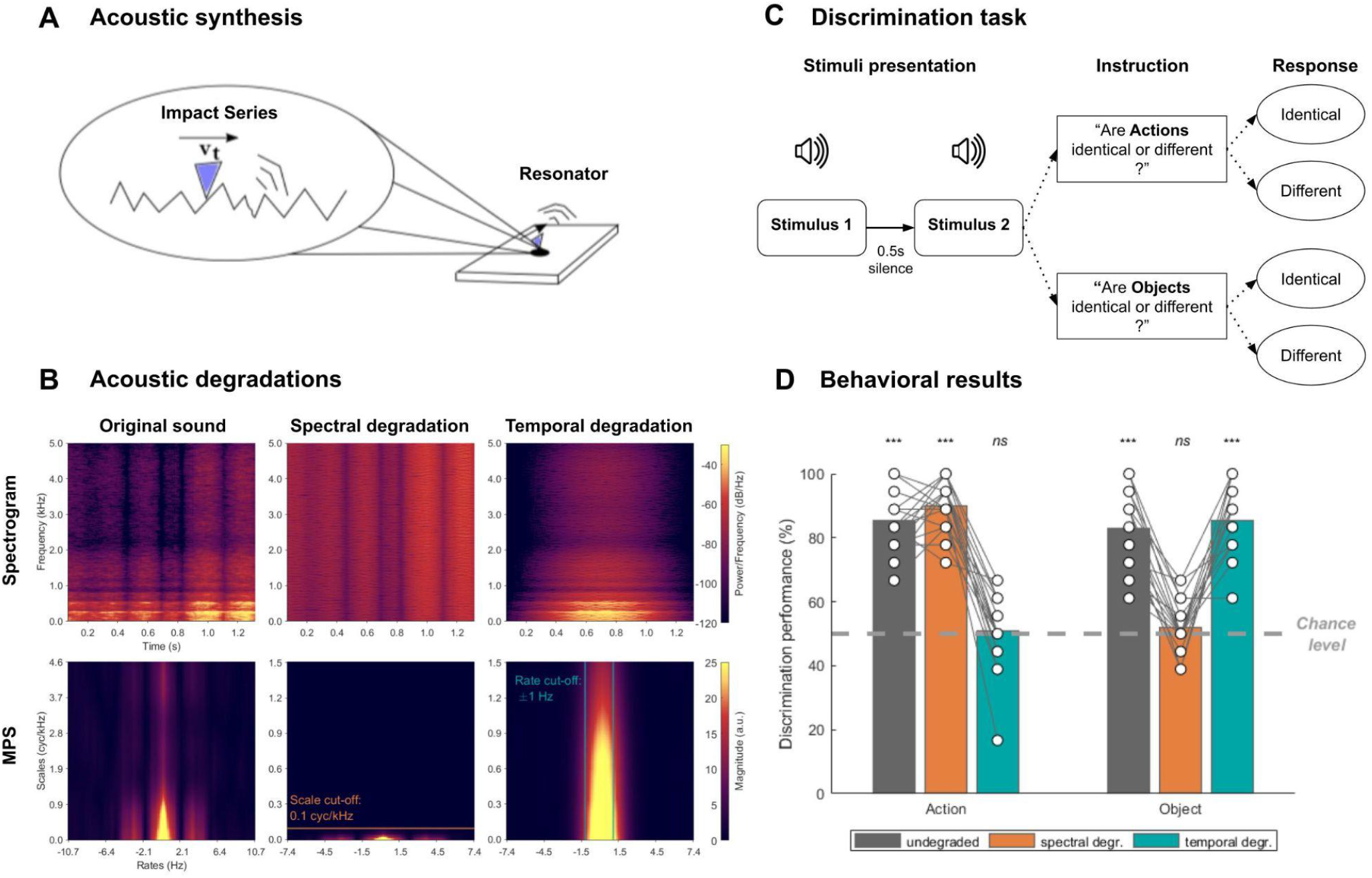
Stimulus set and behavioral experiment. **(A)** Each sound was synthesized with a method emulating a friction as a series of impacts of a resonator on a surface. The friction movement (action) governs the density and strength of impacts, and the surface (object) defines the sound of each single impact. **(B)** Spectrogram (top) and MPS (bottom) of an exemplar original stimulus and of its spectrally and temporally degraded counterparts. **(C)** Paradigm. Participants were presented with pairs of sounds, both undegraded, temporally degraded, or spectrally degraded. They were subsequently instructed to focus on a domain of interest (action or object), and to perform a two-alternative forced-choice discrimination task on this domain. **(D)** Behavioral results. Average performance across participants (bars) and individual performance (white dots). N = 21; *** : adjusted p < 0.001; ns : adjusted p > 0.05 for one-tailed t-tests.

In a subsequent behavioral task, we extended the paradigm developed by Albouy and colleagues (2020) to this set of environmental sounds. Participants (n=21) were presented with pairs of stimuli (both of the same degradation type: undegraded, spectrally or temporally degraded) and were asked to state whether the *actions* or the *objects* were different (Figure 2C). Performance was above chance level in all conditions (all p < 0.001, FDR-corrected), except for action recognition of temporally degraded stimuli, and for object recognition of spectrally degraded stimuli, which were not different from chance (all p > 0.245; Figure 2D). These results show that action discrimination relies solely on purely temporal cues and object discrimination relies solely on purely spectral cues. A significant *dom*ain (action/object) × *degradation type* (no/temporal/spectral) interaction (2 × 3 repeated-measures ANOVA: F(2, 40) = 144.73, p < 0.001) confirmed this double dissociation. No significant main effects of domain (F(1, 20) = 1.53, p = 0.230), nor between spectrally and temporally degraded sounds (p = 0.249) were observed.

### 3. Neural decoding of actions and objects depends on activity patterns in the left and right hemispheres respectively

We then investigated the brain response related to action and object processing in a fMRI experiment. We recorded the blood oxygenation level dependent (BOLD) activity of the same volunteers who had participated in the *action-object* behavioral experiment (n = 20) while they completed a 1-back task to ensure that they listened attentively to the sounds. Univariate analyses highlighted large significant activations in bilateral auditory (bilateral Heschl’s gyrus and STG) and inferior frontal areas (Suppl. Figure S4; p <0.05, FWE-corrected, cluster extent >10). An ANOVA comparing the activity between the three actions or three objects failed to reveal any significant difference (all p > 0.9; FWE-corrected, cluster extent >10) for any of the degradation types.

To investigate more fine-grained encoding of individual actions and objects, we conducted multivariate pattern analyses (MVPA) with a whole-brain searchlight procedure. Specifically, we sought to identify clusters whose response profile is compatible with the general pattern of behavioral performance measured in the first experiment. That is, on the one hand voxels with above-chance decoding accuracy of individual actions for undegraded and spectrally degraded (but not for temporally degraded) stimuli; and on the other hand, voxels with above-chance decoding accuracy of individual objects for undegraded and temporally degraded (but not for spectrally degraded) stimuli (Figure 3A). Such patterns of accuracy results would ensure that the underlying brain activity encodes the purely temporal and purely spectral cues on which actions and objects recognition are based.

**Figure 3:**
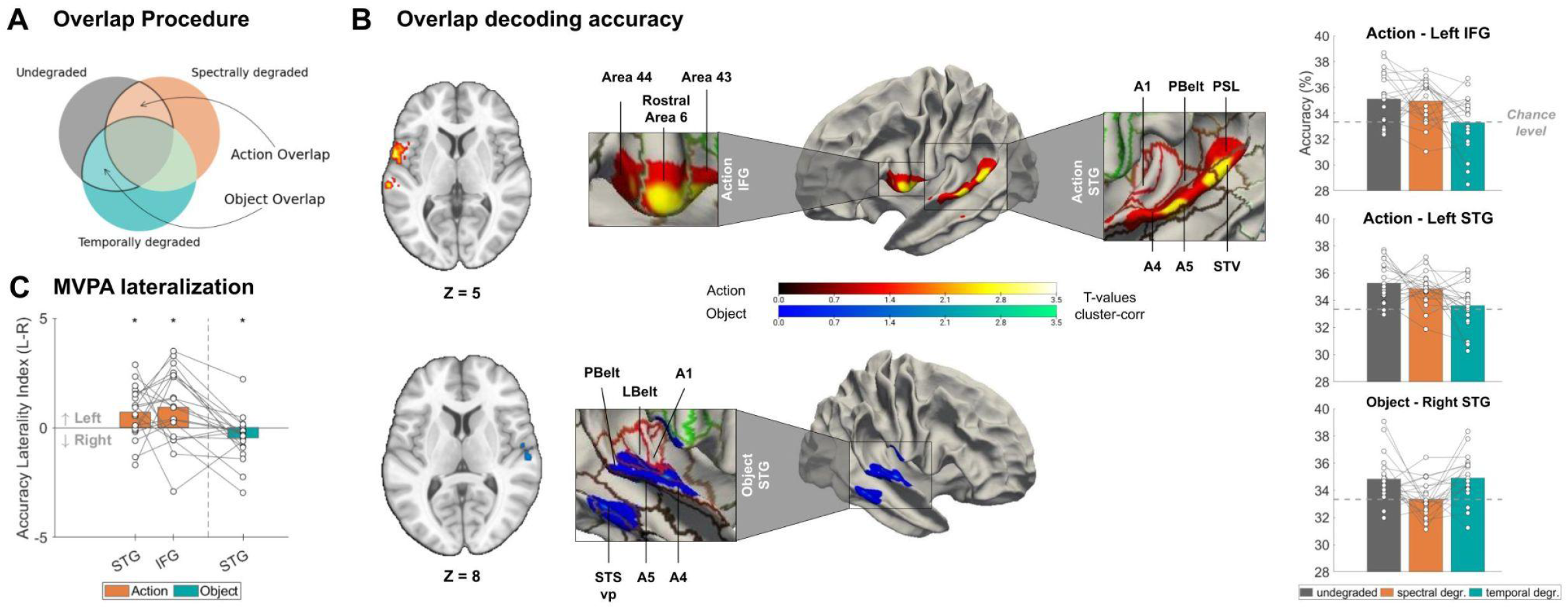
Neural decoding of actions and objects. **(A)** Illustration of the overlap procedure. For each condition, decoding accuracy maps were tested against chance at the group-level and thresholded. The action-overlap map reports significant clusters with above-threshold decoding accuracy of individual actions for undegraded and spectrally degraded (but not for temporally degraded) stimuli. The object-overlap map reports significant clusters with above-threshold decoding accuracy of individual objects for undegraded and temporally degraded (but not for spectrally degraded) stimuli. **(B)** Left Panel: Overlap decoding accuracy maps for 3-class classification of actions and objects (N = 20, p < 0.05, cluster-based permutation test). The T-values displayed correspond to the minimum of the group-level t-values in the two overlapping conditions of interest. Center Panel: T-values plotted on FsAverage surface (gray/white matter boundary). Each zoomed area corresponds to a significant 3D cluster. Regions of interest are extracted from the atlas by Glasser et al. (2016). Right Panel: Decoding accuracy in significant clusters of (B) presented as a function of degradation type (chance level at 33.3%). **(C)** Accuracy laterality index (L-R) in significant clusters, averaged across degradation types. Positive values indicate left-lateralized accuracy, whereas negative values indicate right-lateralized accuracy. Bar plots show mean accuracy. White circles indicate individual data. * : adjusted p < 0.05 for one-tailed t-tests.

Three-class classification (support-vector machine, leave-one-out cross-validation procedure) of either actions or objects was performed separately for each degradation type. Group-level accuracy maps were tested against chance-level (33.3%), yielding statistical parametric maps for each condition (Suppl. Figure S5). These maps were then thresholded and overlapped. In the action-overlap analysis, all above-threshold voxels in undegraded and spectrally degraded conditions were kept, and all above-threshold voxels in the temporally degraded condition were excluded. In the object-overlap analysis, all above-threshold voxels in undegraded and temporally degraded conditions were kept, and all above-threshold voxels in the spectrally degraded condition were excluded (Figure 3A). Cluster-based permutation test revealed that actions were encoded in the left auditory cortex (one cluster encompassing Auditory 4 and 5 complexes (A4/A5); p = 0.002) as well as in the left inferior frontal gyrus (encompassing Area 44 and Rostral Area 6; p = 0.006; see Glasser et al., 2016; Figure 3B). In contrast, objects were encoded in neural patterns of activity in the right auditory cortex (encompassing Parabelt, A4 and A5, ventral posterior Superior Temporal Sulcus; p = 0.036). Similar results were obtained when replicating the overlap procedure without the exclusion of the condition of non-interest (Suppl. Figure S7).

To explicitly test the hypothesis of an hemispheric asymmetry in the encoding of purely temporal and purely spectral modulations, we investigated whether the neural decoding patterns of actions and objects were significantly lateralized. For each cluster, we defined its counterpart in the contralateral hemisphere and computed the mean accuracy lateralization index (L-R) across all degradation types (results per degradation type are displayed in Suppl. Figure S6). We first observed that the three clusters were differently lateralized (one-way repeated-measures ANOVA: F(2, 19) = 7.214, p = .002). We then found a significant leftward dominance for action decoding in the left auditory cortex (p = 0.017) and in the left inferior frontal gyrus (p = 0.017), and a rightward dominance for object decoding in the right auditory cortex (p = 0.030; one-tailed t-tests, FDR-corrected; Figure 3C).

## Discussion

In this study, we investigated whether the action/object framework provides a comprehensive explanation of auditory hemispheric asymmetry at the functional level of Marrian analysis (Marr, 1982). We present three complementary pieces of evidence in favor of this conclusion. First, we show that the acoustic structure of corpora of natural environmental sounds can be well-described by independent combinations of *purely* temporal and *purely* spectral modulations. This indicates that the STM framework is a general framework of auditory processing, encompassing all sounds. Second, we synthesized environmental sounds under the action/object framework to manipulate their temporal and spectral content independently. A behavioral experiment showed that *action* perception relies solely on purely temporal modulations, while *object* perception relies solely on purely spectral modulations. This double dissociation shows that the action/object framework is fully compatible with the STM asymmetry hypothesis, each describing a specific level of analysis. Third, an fMRI experiment revealed that the neural decoding of actions and objects depends on activity patterns in the left and right hemispheres, respectively. This demonstration of an asymmetric processing of action and object provides an account of auditory hemispheric asymmetry in which each hemisphere is specialized for different STM ranges which map onto two categories of invariant acoustic cues - actions and objects - found in all sounds, making it an ecologically valid explanation that aligns with the theory of efficient neural coding (Barlow, 1968; Smith and Lewicki, 2006; Gervain & Geffen, 2019).

The reason why humans can so easily detect and recognize different sounds is because they capitalize on distinctive acoustic features. Recent works have shown that communicative signals (e.g. alarm, emotional, linguistic, song) exploit distinct acoustic niches to target specific neural networks and trigger reactions adapted to the intent of the emitter (Albouy et al., 2020; Arnal et al., 2019; Albouy et al., 2023). Using neurally relevant spectro-temporal representations (the STM framework), these works show that different subspaces of the STM encode distinct information types: temporal modulations for meaning (speech), spectral modulations for melodies, or very fast temporal modulations for alarms (screams). These latter are also critical to discriminate between different soundscapes of natural spaces (Thoret et al., 2020). Here, by analyzing corpora of natural environmental sounds, we show that sounds are mostly composed of purely temporal and purely spectral modulations (Figure 1), at ranges also observed in speech and music. In contrast, combined very fast temporal and high spectral modulations (i.e. their interaction) seem to be mostly visible in alarm sounds, and could reflect an acoustic niche, to drive the alertness network, through non-lemniscal pathways, in essence by-passing classical auditory routes (Arnal et al., 2015, Arnal et al., 2019). Whether such sounds can be described under the action/object framework and show a lateralization profile of response remains to be investigated. Critically, our behavioral results provide clear and strong evidence that temporal and spectral cues directly contribute to the categorization of the two invariant structures composing any sound (Figure 2). While transformational invariants characterize the properties of the action such as its dynamics (Thoret et al., 2014; Conan et al., 2014b), and are encoded in purely temporal modulations, structural invariants characterize the properties of the object such as its surface properties or material (Aramaki et al., 2010; Aramaki et al., 2011) and are encoded in purely spectral modulations.

At the neural level, we reveal that the action/object framework is fully compatible with the STM asymmetry hypothesis (Flinker et al., 2019; Figure 3). Our results replicate and generalize our recent findings showing that a distinct hemispheric sensitivity to STM appears as the mediating mechanism underlying the lateralized responses to speech and melodies (Albouy et al., 2020). We observe significant clusters located in the associative auditory cortex bilaterally (A4), homologous to those of Albouy and colleagues, and show that the STM framework is a general theory of auditory processing, rather than being exclusive to speech and music processing. That lateralization emerges in the associative auditory cortex is in line with a recent study that specifically investigated on a large dataset of intracranial EEG whether asymmetry emerged in the primary, secondary or associative auditory cortex (Giroud et al., 2020). Surprisingly, we also reveal a left-lateralized response for action processing in the inferior frontal gyrus (Area 44, Rostral Area 6), indicating that even asymmetries in the frontal lobe, classically thought to reflect domain-specific processes, such as the left-lateralized language activations (Zatorre and Gandour, 2008; Fedorenko & Blank, 2020) can instead be driven by purely acoustic-specific features. Despite our stimuli containing relatively slow temporal (<5 Hz) and low spectral modulations (<5 cycle/kHz), we highlight a clear hemispheric asymmetry in auditory processing. This finding indicates that asymmetry does not only emerge for extreme STM ranges, but already for the range of STM observed in natural sounds. Moreover, we confirm that while pitch processing does lead to some hemispheric asymmetry (Hyde et al., 2008), it is not the sole driver of lateralized auditory responses. Indeed, we rule out the hypothesis that right-lateralized auditory responses are related to pitch processing, as our stimuli were generated to be non-periodic and devoid of pitch information (Suppl. Figure S3). Instead, our results are in line with the proposal (Schönwiesner et al., 2005; Boemio et al., 2005) that a fine spectral resolution allows for efficient processing of timbre, a key factor in environmental sound perception.

Prior studies have proposed that STM are processed in the auditory cortex along an antero-posterior continuum along the superior temporal gyrus (Schönwiesner & Zatorre, 2009; Santoro et al., 2014). Importantly, our study did not use univariate analyses to map sounds with specific acoustic content, but instead characterized the encoding of cognitive dimensions (action/object) that are derived from acoustic features. Our focus is hence on the functional role of hemispheric asymmetry, beyond acoustic-level processes. We also did not distinguish between general sound categories, such as speech/music or songs, but directly targeted the encoding of individual examples of general cognitive dimensions. This could explain the apparent contradictory findings between recent studies observing (Flinker et al., 2019, Albouy et al., 2020) or failing to observe (Norman-Haignere et al., 2015; Boebinger et al., 2021; Norman-Haignere et al., 2022) auditory hemispheric asymmetry for music and speech. For instance, while selective responses to voices and music *auditory domains* occur bilaterally in associative auditory regions (Norman-Haignere et al., 2015; Boebinger et al., 2021; Norman-Haignere et al., 2022), processing of *individual* sentences and melodies occur in the left and right hemispheres respectively (Albouy et al., 2020). In other words, neural response patterns shared across all stimuli of the same domain (voices, music) are present bilaterally, while neural patterns discriminating different instances of the same domain are more focal and lateralized. This lateralization arguably reflects the complementary specialization of two neural systems functioning in parallel in each hemisphere to maximize the efficiency of the encoding of their respective acoustical features (purely temporal vs. purely spectral modulations). The stages of auditory analysis are sequentially anchored in a cortical processing hierarchy. Starting bilaterally with the rapid identification of the relevant cognitive domain (voice, music), the routing of auditory information obeys a functional division of labor entailing the lateralized specialization of anterior temporal regions for the parallel processing of complex affordances (i.e. meaning, affect and identity). In accordance with this proposal, different auditory clusters are observed between studies investigating general categories (Norman-Haignere et al., 2015) and those decoding individual stimuli (Albouy et al., 2020).

Hemispheric asymmetry is ubiquitous in the animal kingdom, in both vertebrates and invertebrates, and predates humans by half a billion years (Güntürkün & Ocklenburg, 2017; Rogers et al., 2013). In humans, it is most prominent in auditory and somatosensory regions (Geschwind & Galaburda, 1985; Roe et al., 2021). Critically, the most obvious sign that our brains function asymmetrically is the near-universal preference for the right hand (∼9/10 of humans, Papadatou-Pastou et al., 2020). However, even this asymmetry originates in auditory sensory cortices, a result observed during speech production (Kell et al., 2011), hand dynamic actions (Pflug et al., 2019) and speech feedback control (Floegel et al., 2020). Here we propose that such asymmetry can be explained as an adaptation to environmental sounds, with the left hemisphere optimized for processing temporal cues, which are related to actions, while its right counterpart is optimized for processing spectral cues, which are related to objects—a functional distinction allowing the efficient categorization of any environmental sound. We propose that such asymmetry can be the evolutionary solution to the Gabor-Heisenberg uncertainty. The spectro-temporal trade-off in sound analysis is implemented structurally in auditory cortical regions with differential neural populations, resulting in an asymmetry in STM processing which translates into a specialization for action and object processing in the left and right associative auditory regions respectively.

## Materials and Methods

### Analysis of a corpus of environmental sounds

The corpus of environmental sounds is composed of two datasets, the first one from McDermott and Simoncelli (2011) and the second one from Norman-Haignere and colleagues (2015), yielding a total of 256 sounds. Their durations have been cut to 2 seconds to normalize across datasets, and they were down-sampled at 16kHz and normalized to a fixed maximum amplitude.

The modulation power spectrum (MPS) of each sound was computed in MATLAB (Version R2021a) with the Modfilter toolbox (Elliot & Theunissen, 2009), with a windows-size of 166 points and a frame-step of 90 points. Two subspaces of the MPS were defined: the first one corresponds to purely temporal modulations and excludes all spectral modulations above 0.1 cycle/kHz, the second corresponds to purely spectral modulations and excludes all temporal modulations above 土1Hz. The range of values (max scale, min/max rate) of the MPS was chosen so that the overall MPS magnitude across the corpus is identical in each subspace (Figure 1C).

An Independent Component (IC) Analysis with 2 components was performed on the MPSs with the fastICA MATLAB package (https://research.ics.aalto.fi/ica/fastica/, Hyvarinen, 1999).

### Stimuli

#### Stimuli synthesis

The stimuli were generated with MATLAB according to the analysis by synthesis approach (Risset & Wessel, 1999) from which is derived the *action-object* synthesis paradigm (Conan et al., 2014b). This method was designed to synthesize different trajectories of friction motions (actions) on different surfaces (objects), which combines two independent cognitive domains. The working principle of this model consists in the simulation of a series of impacts of an exciter on the asperities of the surface of a resonant object (Figure 2A). From a signal processing point of view, such a principle can be modeled as the convolution between a random signal, characterizing the series of impacts, with an impulse response, characterizing the resonance of the object. Crucially, it allows us to control separately the two domains (action and object) across stimuli, which wouldn’t have been possible with recorded sounds.

Actions are primarily defined by the time-varying velocity of the friction motion. Here, the velocity profile is built upon four oscillators with different frequencies and phases, according to the following equation:

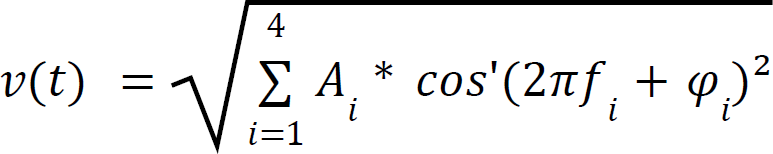

with *f*_1_ 2.5 *Hz* and *f*_2_ to *f_4_* set at random between 0 and 2.5 Hz, phase φ_1_ to φ_4_ set at random, and *A*_1_ = 0. 51 so that the first oscillator does not overwhelm the others (Suppl. Figure S1A). We used this equation to randomly define three different velocity profiles. Each one can be interpreted as erratic friction motions (Thoret et al., 2014). Since *f*_1_ is set at the constant rate of 2.5 Hz and other frequencies are lower, each velocity profile contains modulations at 5 Hz and below. Then, the velocity profiles are used to generate three different series of impacts through a probabilistic model (binomial law), so that a higher velocity yields a larger number of impacts (Suppl. Figure S1B).

Differences between objects were based on surface roughness. Surface roughness can be seen as the interaction between the size of the surface asperities, their depth and their length, and the size of the exciter in contact with the object surface, e.g., the diameter and the length of the pencil nail. Here, we defined three random rugosity profiles (random arrays of 100 samples) to simulate different surface roughness (Suppl. Figure S1C). Then, an impulse response (brief dampened sound, f0 at 260 Hz, 69 other frequencies as multiple of the f0) was generated (Suppl. Figure S1D). The spectral content of the impulse response was kept constant to limit potential pitch differences between sounds. Rugosity profiles were then convolved with the impulse response to yield the modal response of each object (Suppl. Figure S1E). Finally, modal responses were convolved with the impact series (actions) to generate the waveforms (Suppl. Figure S1F). The spectrum of each resulting sound contains peaks at 260, 520 and 760 Hz, and displays broad differences in spectral envelope above 1000 Hz between objects (Suppl. Figure S2).

We ended up with 9 auditory stimuli (available online: https://osf.io/3k78c/), containing all the possible combinations between the 3 velocity profiles (actions) and the 3 surface properties (objects) previously defined (Suppl. Figure S2).

#### Temporal and spectral degradations

Stimuli were degraded on the temporal or spectral dimension with a filtering method developed by Elliott and Theunissen (2009). First, a 2D-Fast Fourier Transform (2D-FFT) is applied on the magnitude of the spectrogram of each sound to estimate their modulation power spectrum (MPS). The resulting MPS represents the temporal (in Hz) and spectral modulations (in cycle/kHz) of a sound. Then, we applied either a 1 Hz cut-off on the temporal dimension or a 0.1 cycle/kHz cut-off on the spectral dimension of the MPSs to remove all modulations above these thresholds.

The filtered MPSs were then transformed back into a spectrogram with an inverse 2D-FFT, and an iterative convex projection algorithm (Griffin & Lim, 1984) was used to recreate the sounds that maximally match the desired spectrogram (Figure 2B). The final set of stimuli thus contains 27 unique stimuli: the 9 undegraded (see above), their 9 temporally degraded counterparts and their 9 spectrally degraded counterparts, all peak-normalized at -10 dB and with a duration of 1.4 seconds.

#### Periodicity analysis

Periodicity distinguishes between sounds with peaks (regularly spaced or not) within their spectrum and non-stationary sounds, such as noise, which have a broad-band spectrum. To evaluate the periodicity of our stimuli, we looked at the maximum of their time-varying cepstrum. The cepstrum is obtained by performing a fourier transform on the time-varying power spectrum of a sound, and its magnitude at a given quefrency indicates the presence of frequencies that are multiples of a common f0 - even when the f0 is not present in the sound. Cepstrums are notably used in pitch determination algorithms (Noll, 1967). The initial fourier transform was computed with a maximal frequency of 10kHz, yielding a cepstrum frequency range of 100 to 5000 Hz, corresponding to the range of pitch percepts (Attneave and Olson, 1971; Pressnitzer et al., 2001). To compare our stimuli with sounds that elicited rightward dominance in a previous study, we also applied this periodicity measure to the *a capella* songs from Albouy and colleagues (2020).

### Behavioral experiment

#### Participants

21 (20+1 excluded, see below) healthy young adults (15 females, mean age = 23.3 ± 3.7 years) participated in an online experiment hosted on FindingFive (https://www.findingfive.com), all right-handed, with normal hearing and no known neurological disorders. No participants had more than 5 years of recent musical practice. Informed consent was obtained for all participants. The experiment was granted ethical approval by Aix-Marseille Université “Comité Protection des Personnes”—“Committee for the Protection of Individuals”: CPP 2017-A03614–49. They were given 40 euros for their participation (for both the online and fMRI experiments)

#### Procedure

All participants completed an online discrimination task hosted on FindingFive. They were asked to complete it in a calm environment, with earphones or headphones. The experiment lasted approximately 25 minutes. In each trial, a pair of stimuli was presented, separated with 0.5 second of silence. After the stimulus presentation, participants were instructed to focus either on the “friction pattern” (*action*) or on the “*object”*, and were asked to state whether the two sounds were identical or different on the relevant domain (Figure 2C). The pairs of stimuli were identical for half of the trials (chance level = 50%) on the domain of interest (*action* or *object*), always different on the domain of non-interest, and the two sounds of a given pair always belonged to the same degradation type (undegraded, temporally degraded, or spectrally degraded).

At the beginning of the experiment, four examples of friction patterns and objects were provided (the “familiarization stimuli”, different from the stimuli used for the actual experiment), to ensure that the participants successfully understood the task and to allow them to set the sound volume at a comfortable level. Then, they completed a familiarization session on 16 pairs of undegraded sounds (training stimuli). After the training phase, a few examples of temporally and spectrally degraded sounds (training stimuli) were provided, so that participants were familiarized with these types of degraded sounds prior to the actual experiment. Afterwards, they completed 3 blocks of 36 trials each (test stimuli), with all degradation types (undegraded, temporal, spectral), instruction (focus on action or object) and expected answer (same/different) equally balanced and randomly ordered.

#### Statistical analysis of behavioral data

Statistical analyses were conducted on MATLAB. They were performed with the percentage of correct discrimination as the dependent variable. One-tailed t-tests were conducted on each condition separately (with Holm-Bonferroni correction) to test whether the performance accuracy was above the chance-level (Figure 2D). A two-way repeated measures ANOVA was conducted with the two within-subject factors domain (action/object) and degradation type (undegraded, temporal, spectral) to investigate main effects and interactions, and post-hoc tests were computed with Holm-Bonferroni correction for multiple comparisons.

### FMRI experiment

#### Participants

The participants in the fMRI experiment were the same ones who had already completed the online behavioral task. One participant moved too much during the fMRI experiment, making their data unworkable, and has been removed from the data set. The data used for the subsequent analyses thus contain 20 participants.

#### Procedure

The experiment lasted 41 minutes, divided in 6 runs of 6-7 minutes each. In each run, the 27 unique stimuli were presented twice in total, in random order, with a jittered interstimulus interval of 4, 6 or 8 seconds. 12 stimuli were randomly selected and presented two times consecutively. The second presentation is the catch trial for the 1-back task they had to perform (see below). A 20 second break was introduced in the middle and at the end of each run. Overall, the entire stimuli set was presented 12 times, and there were 72 catch trials across the whole experiment. Sensimetrics MRI-compatible insert earphones were used to display the sounds to the participants (∼90 dB). Stimuli presentation was done using a specific software using the NI LabVIEW 2017 programming environment and a NI PCIe-6536B (National Instruments, Austin, Texas, U.S.) digital card in order to synchronize stimuli diffusion with MRI acquisition.

Participants completed a 1-back task, to ensure that their attention was focused on the stimuli throughout the experiment. They were asked to press the response button when they heard a stimulus that is exactly the same as the previous one (catch trial). Since two sounds in the stimuli set could differ in terms of temporal content, spectral content or degradation type, this 1-back task prompted the participants to focus their attention on all the components of the sounds. On average, they successfully detected 79 ± 15% of the catch trials.

#### fMRI acquisition parameters

The fMRI experiment was carried out at the Center IRM-INT@CERIMED (INT, UMR 7289, AMU-CNRS) installed in the CERIMED (“Centre Européen de Recherche en Imagerie Médicale”, Marseille) with a 3T MRI Scanner (MAGNETOM Prisma, Siemens AG, Erlangen, Germany). 3D MPRAGE (Magnetization Prepared - RApid Gradient Echo) T1-weighted images were acquired with 3-fold GRAPPA acceleration (time to acquisition (TA) = 3:52min; time to repetition (TR) = 2300 ms; time to echo (TE) = 2.98 ms; time to inversion (TI) = 925 ms; flip angle (FA) = 9°; readout in superior-inferior direction; matrix size = 240 × 192 × 256; field of view (FOV) = 240 × 192 × 256 mm3; voxel size = 1 × 1 × 1 mm3).

Functional volumes were acquired with a fast event-related sparse-sampling design, to limit the scanner noise coming from the fast adjustment of magnetic gradients and allow the participants to hear the stimuli at ecological volume (Perrachione & Ghosh, 2013). Here, we had 2 second-long TRs, during which we acquired one functional volume with a TA of 539 ms and left 1461 ms of silence, and the stimuli were always presented 31 ms after the TA (570 ms after the beginning of the TR) so that they could be played during the silent periods.

Functional volumes were acquired with the following parameters: 30 oblique slices aligned along the sylvian fissure, with descending, multiband acquisition (acceleration factor 5), TE = 35.8 ms, FA = 78°; 1.8 mm slice thickness; 0.216 gap; matrix size = 100 × 100; FOV = 200 × 200 mm²; voxel size = 2 × 2 × 2.016 mm3. For susceptibility distortion correction, Spin Echo EPI images with reversed phase encoding directions were collected with similar coverage and the following parameters: TR = 7106 ms, TE= 63.2 ms, FA: 90/180°.

#### fMRI preprocessing

Results included in this section come from preprocessing performed using *fMRIPrep* 20.2.1 (Esteban et al., 2019; Esteban et al., 2018; RRID:SCR_016216), which is based on *Nipype* 1.5.1 (Gorgolewski et al., 2011; Gorgolewski et al., 2018; RRID:SCR_002502).

For each subject, the T1-weighted (T1w) image was corrected for intensity non-uniformity (INU) with N4BiasFieldCorrection (Tustison et al., 2010), distributed with ANTs 2.3.3 (Avants et al., 2008, RRID:SCR_004757), and used as T1w-reference throughout the workflow. The T1w-reference was then skull-stripped with a *Nipype* implementation of the antsBrainExtraction.sh workflow (from ANTs), using OASIS30ANTs as target template. Brain tissue segmentation of cerebrospinal fluid (CSF), white-matter (WM) and gray-matter (GM) was performed on the brain-extracted T1w using fast (FSL 5.0.9, RRID:SCR_002823; Zhang et al., 2001). Volume-based spatial normalization to a standard space (MNI152NLin2009cAsym) was performed through nonlinear registration with antsRegistration (ANTs 2.3.3), using brain-extracted versions of both T1w reference and the T1w template. The following template was selected for spatial normalization: *ICBM 152 Nonlinear Asymmetrical template version 2009c* [Fonov et al., 2009; RRID:SCR_008796; TemplateFlow ID: MNI152NLin2009cAsym]. A similar spatial normalization was performed to a symmetrical standard space (*ICBM 152 Nonlinear Symmetrical template version 2009c [Fonov et al., 2009; RRID:SCR_008796; TemplateFlow ID: MNI152NLin2009cSym]*) for subsequent lateralization analysis.

For each of the 6 BOLD runs, the following preprocessing was performed. First, a reference volume and its skull-stripped version were generated by aligning and averaging 1 single-band reference (SBRefs). A B0-nonuniformity map (or *fieldmap*) was estimated based on two echo-planar imaging (EPI) references with opposing phase-encoding directions, with 3dQwarp (Cox & Hyde, 1997; AFNI 20160207). Based on the estimated susceptibility distortion, a corrected EPI (echo-planar imaging) reference was calculated for a more accurate co-registration with the anatomical reference. The BOLD reference was then co-registered to the T1w reference using flirt (FSL 5.0.9; Jenkinson & Smith, 2001) with the boundary-based registration (Greve and Fischl, 2009) cost-function. Co-registration was configured with nine degrees of freedom to account for distortions remaining in the BOLD reference. Head-motion parameters with respect to the BOLD reference (transformation matrices, and six corresponding rotation and translation parameters) are estimated before any spatiotemporal filtering using mcflirt (FSL 5.0.9; Jenkinson et al., 2002). The BOLD time-series were resampled onto their original, native space by applying a single, composite transform to correct for head-motion and susceptibility distortions. These resampled BOLD time-series will be referred to as *preprocessed BOLD in original space*, or just *preprocessed BOLD*. The BOLD time-series were also resampled into standard space, generating a *preprocessed BOLD run in MNI152NLin2009cAsym space*. Several confounding time-series were calculated based on the *preprocessed BOLD*: a set of physiological regressors were extracted to allow for component-based noise correction (*CompCor*; Behzadi et al., 2007). Principal components are estimated after high-pass filtering the *preprocessed BOLD* time-series (using a discrete cosine filter with 128s cut-off) for the anatomical *CompCor* (aCompCor). Two probabilistic masks (CSF and WM) are generated in anatomical space. The implementation differs from that of Behzadi and colleagues in that instead of eroding the masks by 2 pixels on BOLD space, the aCompCor masks are subtracted from a mask of pixels that likely contain a volume fraction of GM. This mask is obtained by thresholding the corresponding partial volume map at 0.05, and it ensures components are not extracted from voxels containing a minimal fraction of GM. Finally, these masks are resampled into BOLD space and binarized by thresholding at 0.99 (as in the original implementation). Components are also calculated separately within the WM and CSF masks. For the CompCor decomposition, the *k* components with the largest singular values are retained, such that the retained components’ time series are sufficient to explain 50 percent of variance across the nuisance mask. The remaining components are dropped from consideration. The head-motion estimates calculated in the correction step were also placed within the corresponding confounds file. The confound time series derived from head motion estimates were expanded with the inclusion of temporal derivatives for each (Satterthwaite et al., 2013).

#### fMRI univariate analysis

Univariate analyses were conducted at the whole-brain level with SPM12 (Wellcome Trust Centre for Neuroimaging, London, UK) on BOLD data in *MNI152NLin2009cAsym* space.

Preprocessed data were first spatially smoothed with a gaussian kernel (8 mm full-width at half-maximum (FWHM)). A canonical hemodynamic response function (HRF) was used to model the BOLD response to undegraded, temporally degraded, or spectrally degraded sounds separately, using a general linear model (GLM) approach. Further regressors have been added to the model to take into account the BOLD response to catch trials (one per degradation type) and the motor activation occurring when the participants pressed the response button. Additionally, twenty-two nuisance regressors computed through the *fMRIPrep* pipeline were included, with estimates of head motion (translations and rotations on the three spatial dimensions and their derivative) and physiological noise (the five aCompCor explaining the more variance in the WM mask, and the fives ones from the CSF mask).

At the group level, a one-tailed t-test with family-wise error (FWE) correction for multiple comparisons was used on whole brain data to highlight the voxels with significant positive response to the stimuli of interest (all except catch trials).

To investigate differences in brain response between the three classes of action and objects, a one-way repeated-measures ANOVA was performed for each domain (action/object) and degradation type (undegraded, temporally or spectrally degraded) separately. This analysis did not reveal any significant voxels (all p > 0.9, FWE-corrected, cluster extent > 10).

#### fMRI multivariate analysis

Multivariate pattern analyses were conducted on preprocessed BOLD data in subject space (T1w space). BOLD data were first smoothed with a gaussian kernel (2 mm FWHM) with SPM12. Next, the shape of the HRF was estimated for each voxel of each participant individually, given that it has been shown to improve decoding analyses (Pedregosa et al., 2013), and that hemodynamic responses in the auditory cortex peak earlier than the standard canonical HRF included in SPM (Belin et al., 1999). HRF estimation was conducted in a Python environment with the *hrf_estimation* toolbox (Pedregosa et al., 2015). Practically, a rank-one GLM informed with the same design matrix as for univariate analyses (except the button-press events) was used to estimate the shape of the response of each voxel to all auditory events of the experiment (regardless of the experimental conditions), expressed as a weighted sum of a canonical HRF, its derivative and its second derivative. All estimated HRFs were peak-normalized. Then, the response to each trial was estimated with a classical GLM approach, with one regressor per stimulus (324 throughout the entire experiment). Other regressors were used to account for the response to catch trials and to include the nuisance regressors, as in univariate analyses. This step was done for each voxel of each participant separately with the HRF previously estimated for this particular voxel, resulting in one “activation” map (beta map) per trial and per participant.

Classification analyses (decoding) were conducted on the activation maps with a similar procedure to the one used in Albouy and colleagues (2020). They were performed with The Decoding Toolbox (Hebart et al., 2015) in MATLAB and a support vector classifier (SVC) as implemented in LibSVM (Chang & Lin, 2011).

The classifier was trained and used with the data corresponding to each degradation type separately (undegraded, temporally and spectrally degraded, 108 trials each) to perform a 3-class classification (chance level = 33.3%) on either the *action* or the *object* domain (e.g., the classifier had to retrieve whether the presented stimulus contained *action 1*, *action 2* or *action 3* for a given trial, based on the corresponding activation maps). A searchlight approach was used to address the high-spatial dimensionality issue of the fMRI data (sphere radius = 8 mm) and the classifier was used with default parameters (linear SVC, linear kernel, cost parameter = 1). The performance of the classification was assessed through a six-fold leave-one-out cross-validation procedure. Resulting accuracy maps were spatially normalized to the MNI152NLin2009cAsym space (applying ANTs spatial transformations calculated through fmriprep) and smoothed (gaussian kernel, 2 mm FWHM). Group-level analyses were then conducted in SPM12. A one-tailed t-test was performed on accuracy maps to highlight the voxels in which the decoding accuracy was significantly above chance. This procedure was conducted on each condition separately.

#### fMRI overlap analysis

To highlight the clusters in which the pattern of decoding accuracy is similar to the behavioral pattern of results, statistical maps (thresholded at t = 1.328) were overlapped across degradation types and within each domain. That is, in the action-overlap analysis, all above-threshold voxels in undegraded and spectrally degraded conditions were kept, and all above-threshold voxels in the temporally degraded condition were excluded. In the object-overlap analysis, all above-threshold voxels in undegraded and temporally degraded conditions were kept, and all above-threshold voxels in the spectrally degraded condition were excluded (Figure 3A). In the resulting maps, each voxel value represents the minimum of the t-value of the two conditions of interest (undegraded and spectrally degraded for *action*, undegraded and temporally degraded for *object*) obtained at the group level in this particular voxel.

Statistical significance was evaluated through cluster-based permutation tests. Permutation tests were conducted on cluster-size (number of contiguous voxels) to assess the significance of the results. Classification analyses were repeated 500 times for each participant and condition with shuffled class-labels, yielding chance-level accuracy maps. In each permutation, the overlap procedure was reproduced on these maps and the size of the largest cluster was recorded. To correct for multiple comparisons across conditions, only the largest cluster among both the action-overlap and the object-overlap was selected at each permutation. This procedure yielded a distribution of cluster-size under the null-hypothesis. In non-permuted data, any cluster size larger than 95% of the clusters obtained from random permutations is considered significant at p < 0.05 cluster-corrected (Figure 3B). We restricted the number of permutations to 500 to limit the computational cost of this procedure, which allows us to evaluate significance with a minimum p-value of 0.002.

#### Regions-of-interest analysis

The clusters obtained from the overlap analyses were used as masks on ‘raw’ accuracy maps (not smoothed) to extract the mean accuracy per condition (Figure 3B).

#### Lateralization analysis

The lateralization of decoding results was assessed by computing the laterality index (left minus right) for each condition. Accuracy maps (not smoothed) from the decoding analysis were first spatially normalized to a symmetric template (using the normalization procedure from FMRIPREP with ANTs and (MNI152NLin2009cSym as the output space) to allow the comparison between homologous brain areas, and accuracy values from the right hemisphere were subtracted from the homologous values extracted from the left hemisphere. Clusters obtained from the overlap procedure (see above) were also transformed to the MNI152NLin2009cSym space and used as masks. This allowed us to compute the mean accuracy lateralization index across voxels in each cluster (Figure 3C; Suppl. Figure S6). Statistical significance was assessed with one-tailed t-tests, according to the hypothesis that action accuracy is left-lateralized (right-tailed t-test) and object accuracy is right-lateralized (left-tailed t-test). P-values were adjusted with Holm-Bonferroni correction for multiple comparisons.

## Conflict of interests

The authors declare no competing interests.

### Acknowledgments

We thank Philippe Albouy, Bruno Nazarian, the ILCB and the Centre IRM-INT@CERIMED for their scientific support, and the participants for their contribution to this project. Centre de Calcul Intensif d’Aix-Marseille is acknowledged for granting access to its high-performance computing resources.

## Author contributions

P.R, P.B., E.T and B.M. designed the experiment. P.R. and A.G. acquired data. P.R., J.S., and JL.A. analyzed data. P.R, R.Z., E.T and B.M. wrote the paper.

## Supplementary materials

**Figure S1:**
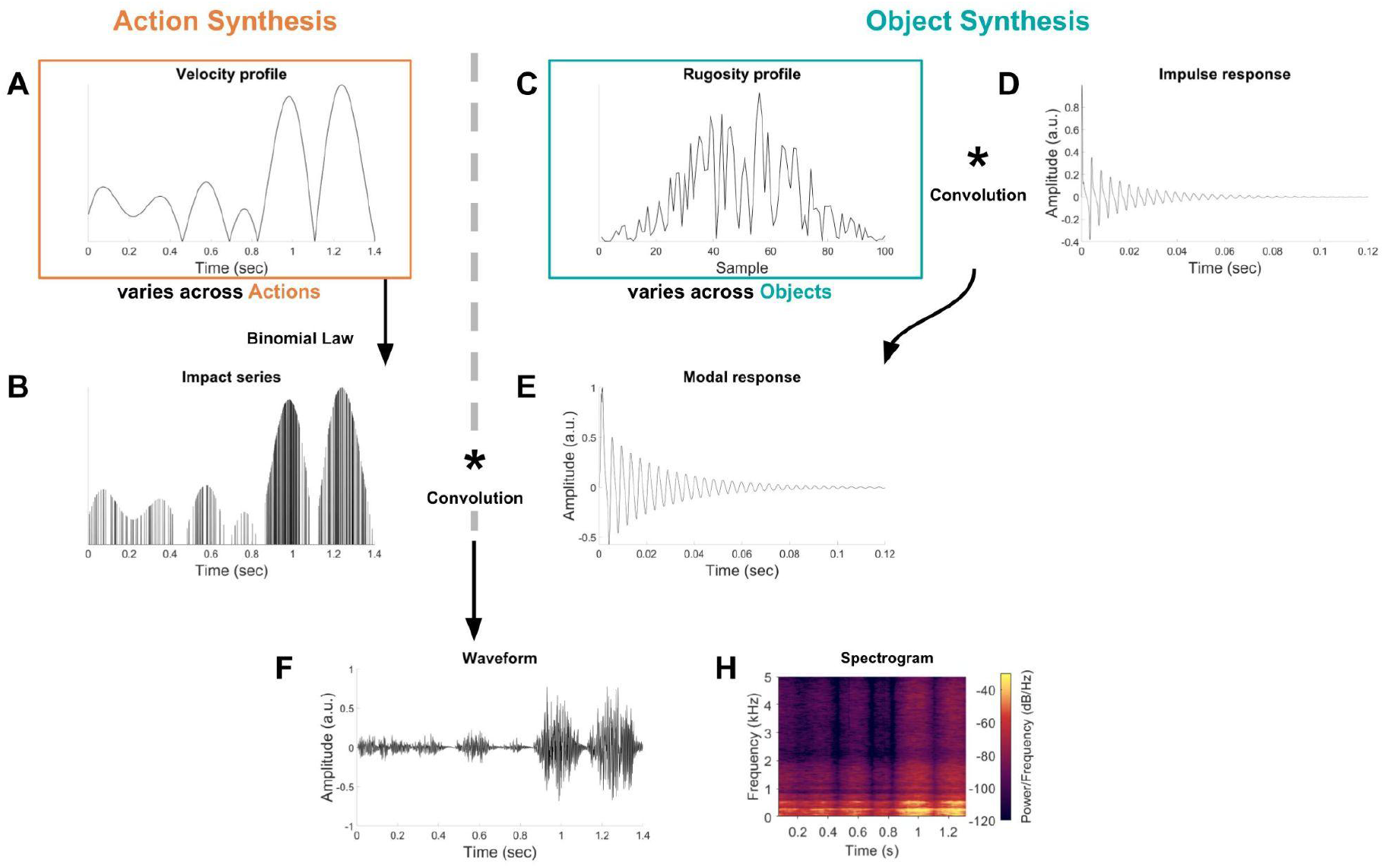
Stimuli synthesis. (A) The velocity profile of a friction movement is built upon several oscillators with varying phase and frequency. These velocity profiles differ across actions. (B) For each action, the velocity profile is converted in a series of impacts through a binomial law. In this figure, impact probability was divided by 10 for visual clarity. (C) An array is generated at random to define the rugosity profile of the friction sound. These rugosity profiles differ across objects. (D) Impulse response defining the sound of a single impact. (E) Rugosity profiles are convolved with the impulse response to produce the modal response of each object. (F) Impact patterns are convolved with the modal response to yield the waveforms. A high-pass filter (threshold = 20Hz) is used to remove slow fluctuations in the final waveforms. (H) Spectrogram of the waveform displayed in panel F.

**Figure S2:**
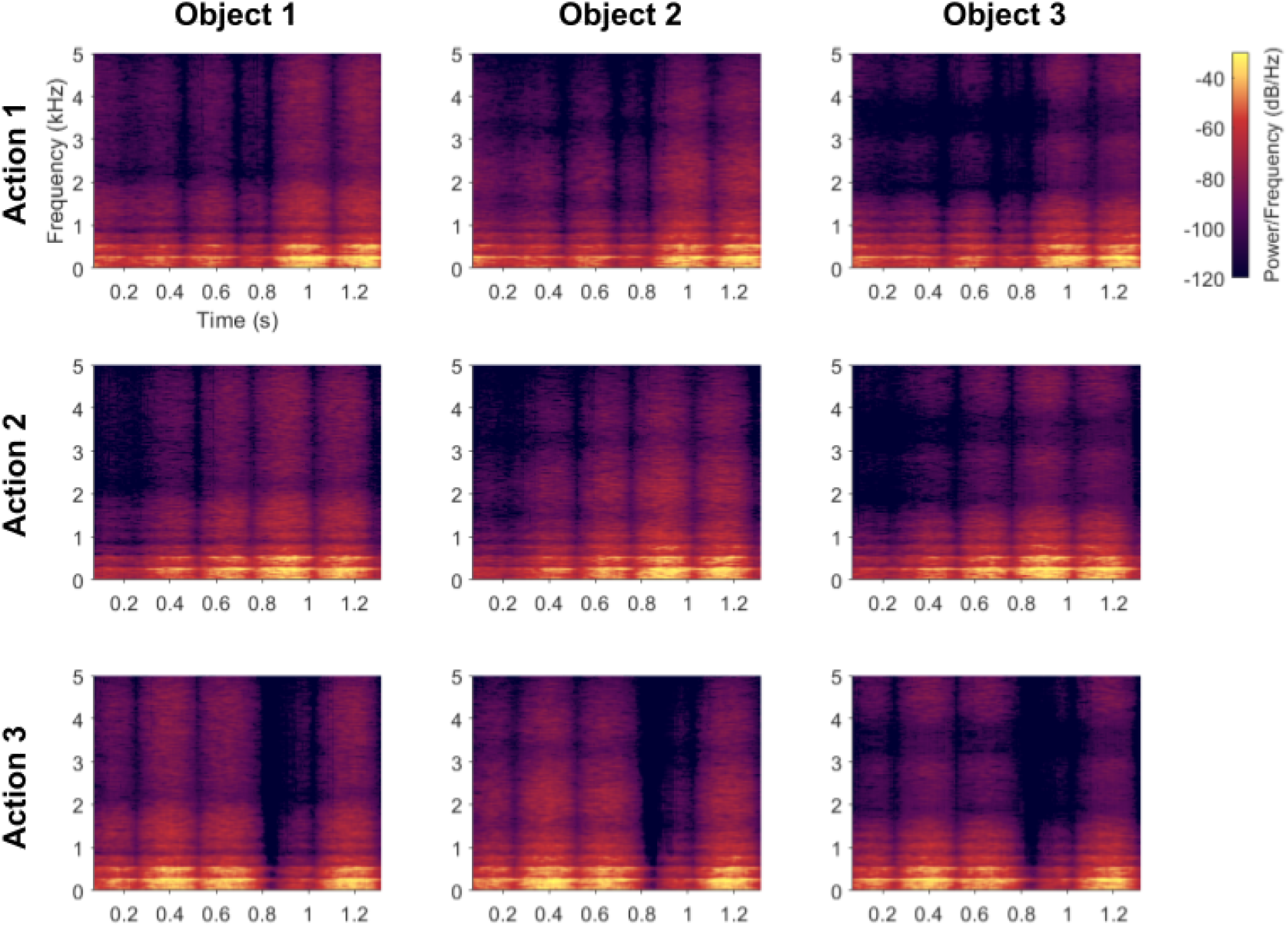
Spectrograms of the 9 undegraded stimuli.

**Figure S3:**
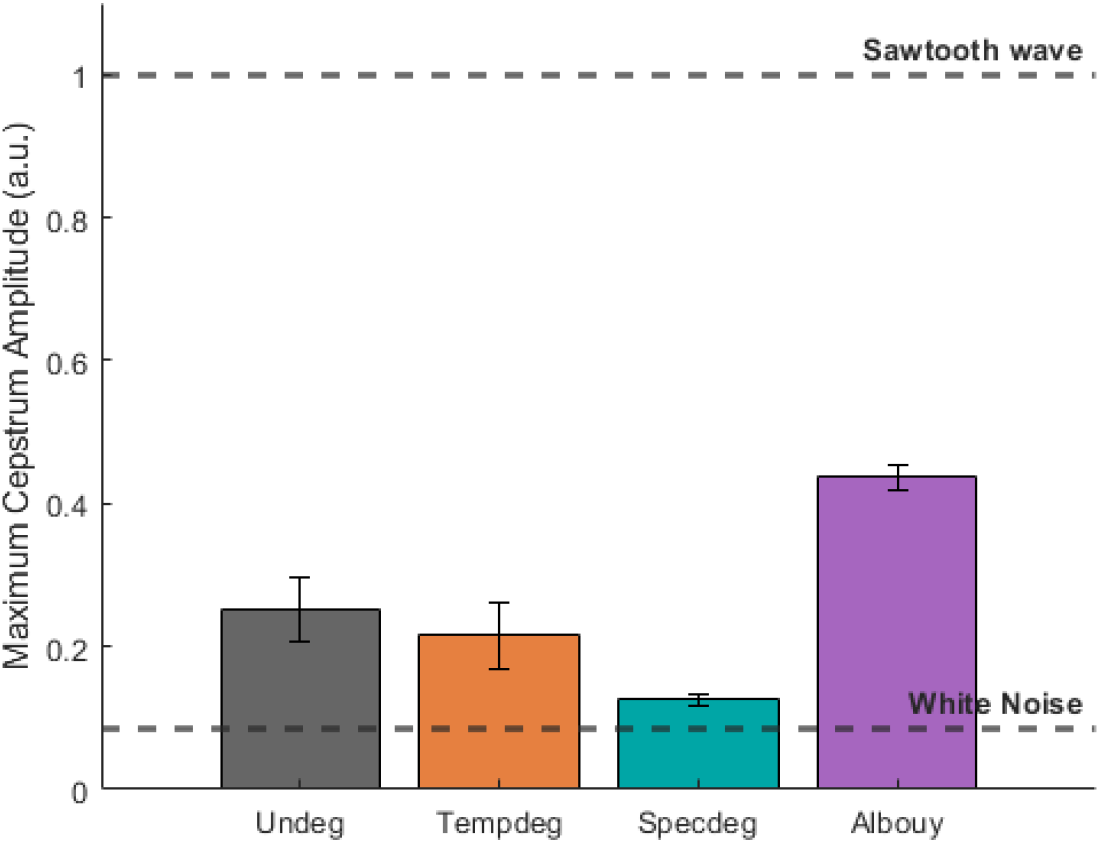
Stimuli periodicity. Maximum of the time-varying cepstrum averaged across stimuli of the same condition. Undegraded (N = 9), temporally (N = 9) and spectrally (N = 9) degraded conditions are depicted, together with the undegraded a capella songs (N = 100) used by Albouy and colleagues (2020). The two extremes of the metric are illustrated by a perfectly periodic sawtooth wave and a non-periodic white noise.

**Figure S4:**
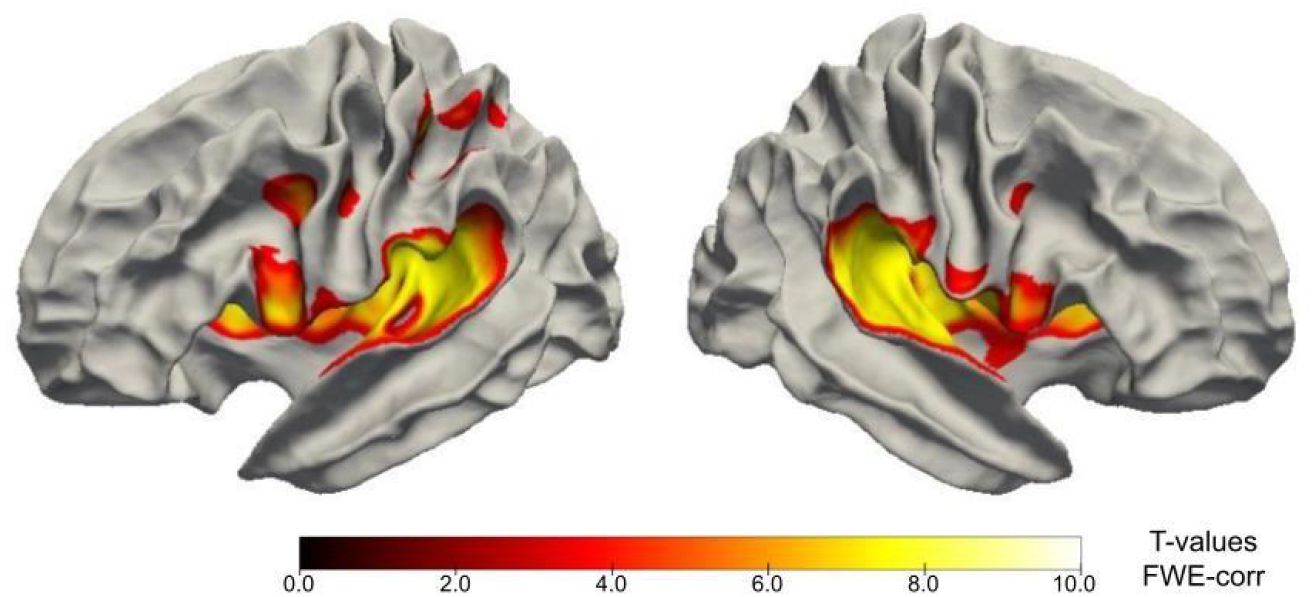
General auditory activation. Significant positive activation in response to all stimuli of interest (N= 20, p <0.05, FWE-corrected).

**Figure S5:**
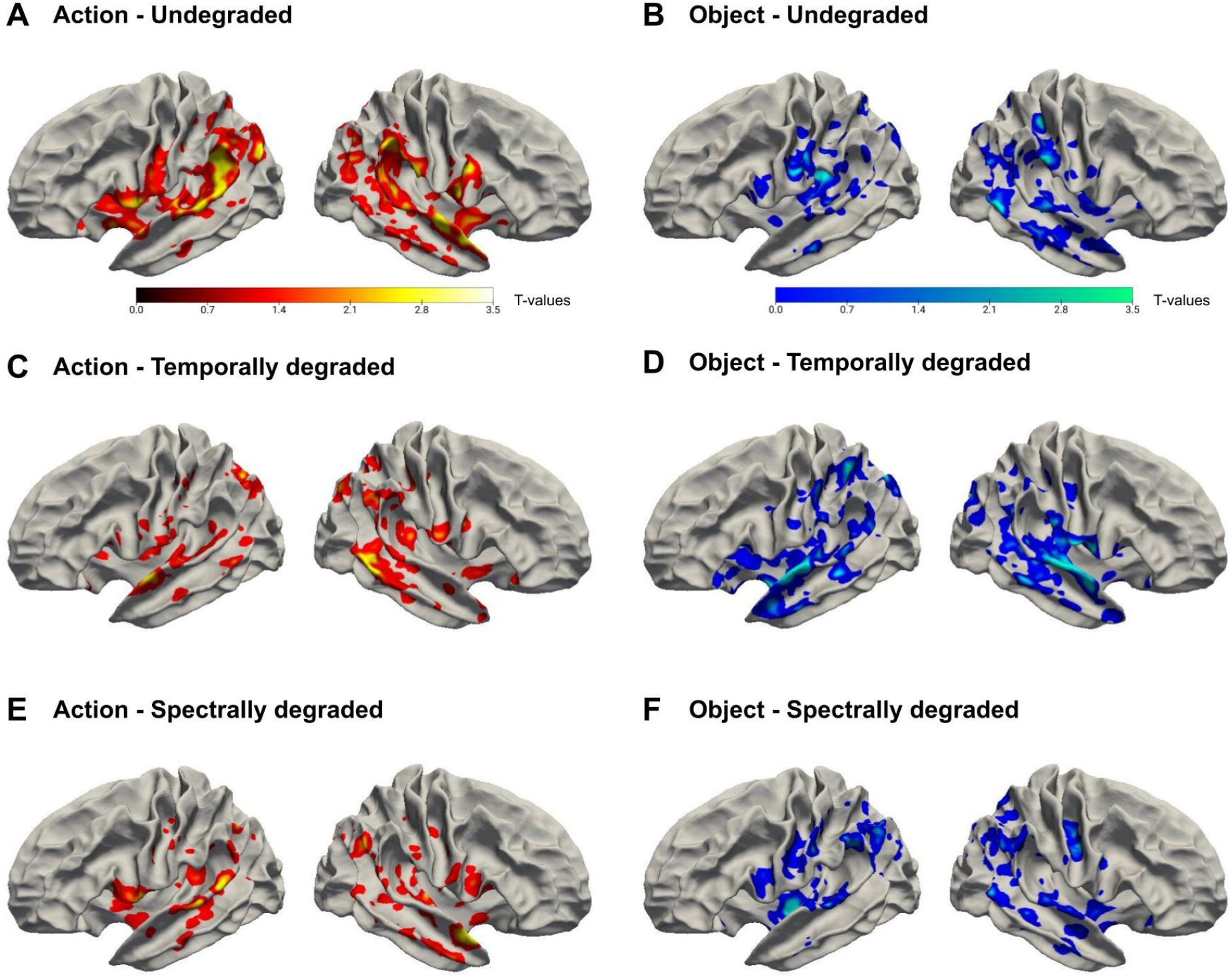
Uncorrected group-level decoding results per condition. Decoding of actions (A, C, E) and objects (B, D, F) for each degradation type (Undegraded: A, B; Temporally degraded: C, D; Spectrally degraded: E, F). T-values (N= 20, p<0.1, uncorrected) are represented with hot (actions) and cold (objects) colors.

**Figure S6:**
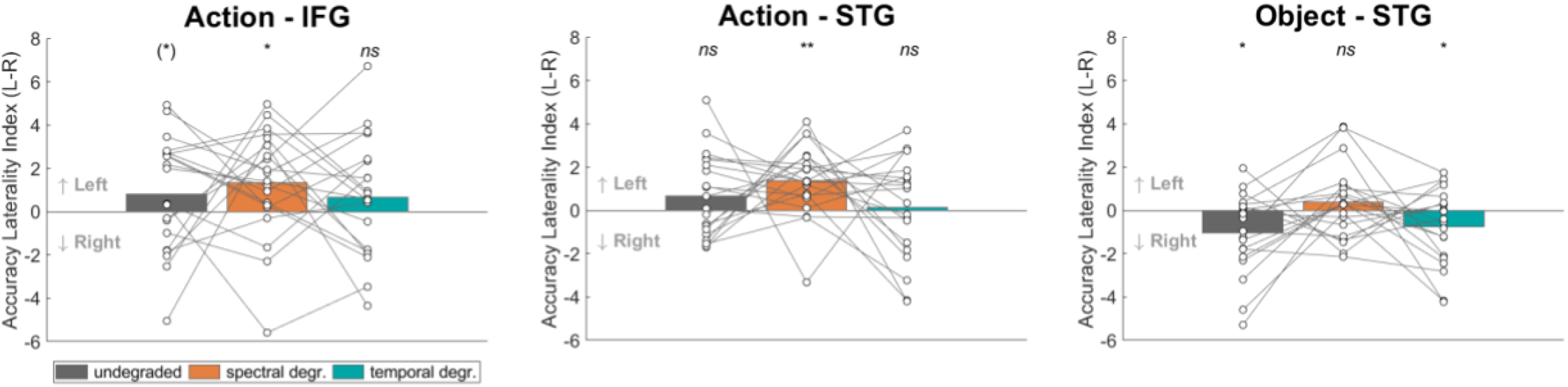
Laterality index per degradation type. Accuracy laterality index (L-R) in significant clusters. Positive values indicate left-lateralized accuracy, whereas negative values indicate right-lateralized accuracy. Bar plots show mean accuracy. White circles indicate individual data (N= 20). ** : adjusted p < 0.01; * : adjusted p < 0.05; (*) : adjusted p < 0.1; ns : adjusted p > 0.1 for one-tailed t-tests.

**Figure S7:**
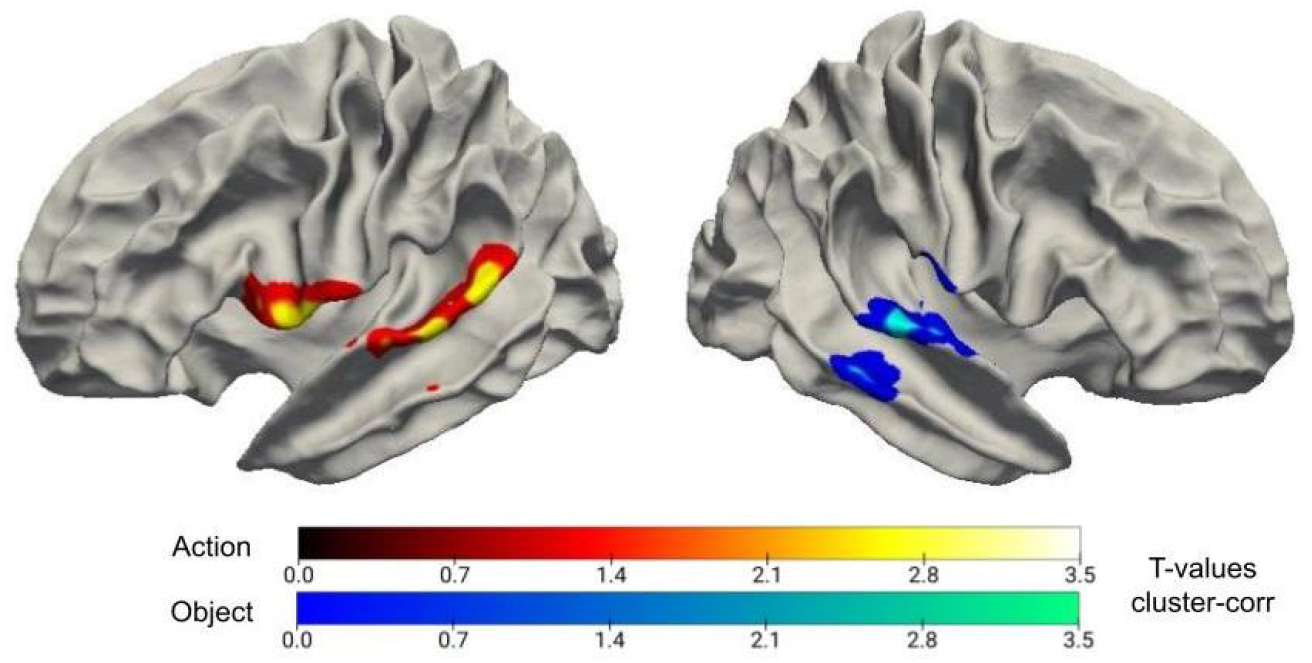
Overlap decoding accuracy maps without exclusion. Action(/object)-overlap with above-threshold decoding accuracy of individual actions for undegraded and spectrally (/temporally) degraded stimuli, without exclusion of above-threshold temporally (/spectrally) degraded ones. Clusters displayed are significant (p < 0.05, permutation test). The T-values displayed correspond to the minimum of the group-level t-values in the two overlapping conditions of interest.

**Table S8:** Summary statistics for each cluster.

| Cluster | Overlap p-value | Cluster size (voxels) | Overlap without exclusion p-value | Peak position (mm) | Minimum peak t-value | Condition | Mean t-value | Maximum t-value | Mean accuracy (%>chance) | STD accuracy | Mean laterality index | STD laterality index |
| --- | --- | --- | --- | --- | --- | --- | --- | --- | --- | --- | --- | --- |
| **Action - Left STG** | **0.002** | 311 | 0.008 | -64.5 ;<br>-38.5 ;<br>17.5 | 4.007 | Overall |  |  | 1.2235 | 0.8788 | 0.7222 | 1.1694 |
|  |  |  |  |  |  | undeg | 2.6163 | 5.5452 | 1.9104 | 1.4649 | 0.6546 | 1.9396 |
|  |  |  |  |  |  | tempdeg | 0.3799 | 1.3213 | 0.2573 | 1.6712 | 0.1362 | 2.3484 |
|  |  |  |  |  |  | specdeg | 1.9827 | 4.0070 | 1.5029 | 1.2178 | 1.3759 | 1.6509 |
| **Action - Left IFG** | **0.006** | 299 | 0.008 | -58.5 ;<br>7.5 ;<br>1.5 | 3.368 | Overall |  |  | 1.1161 | 1.3062 | 0.9395 | 1.6903 |
|  |  |  |  |  |  | undeg | 2.2648 | 4.8415 | 1.7774 | 2.0685 | 0.8159 | 2.6422 |
|  |  |  |  |  |  | tempdeg | -0.0152 | 1.3138 | -0.0529 | 2.1561 | 0.6711 | 2.7495 |
|  |  |  |  |  |  | specdeg | 2.1561 | 5.4821 | 1.6238 | 1.7087 | 1.3315 | 2.5208 |
| **Object - Right STG** | **0.036** | 232 | 0.008 | 59.5 ;<br>-16.5 ;<br>5.5 | 2.411 | Overall |  |  | 1.0289 | 0.8134 | -0.4687 | 1.0472 |
|  |  |  |  |  |  | undeg | 1.9500 | 3.5633 | 1.5015 | 1.7443 | -1.0175 | 1.8498 |
|  |  |  |  |  |  | tempdeg | 1.8732 | 3.7969 | 1.5594 | 1.7164 | -0.7567 | 1.7781 |
|  |  |  |  |  |  | specdeg | 0.0715 | 1.3262 | 0.0258 | 1.3571 | 0.3682 | 1.7005 |
| Action - Right anterior STG | 0.052 | 225 | 0.05 | 55.5 ;<br>9.5 ;<br>-16.5 | 3.542 | Overall |  |  | 1.1592 | 0.9667 | -0.7749 | 1.3373 |
|  |  |  |  |  |  | undeg | 2.3485 | 4.3751 | 1.6536 | 1.4816 | -1.1496 | 2.0258 |
|  |  |  |  |  |  | tempdeg | 0.2796 | 1.3097 | 0.2081 | 1.9699 | 0.2720 | 2.6378 |
|  |  |  |  |  |  | specdeg | 2.2625 | 4.9528 | 1.6159 | 1.6861 | -1.4471 | 2.1754 |
| Object - Left Insula/A1 | 0.1 | 188 | 0.126 | -40.5 ;<br>-22.5 ;<br>11.5 | 2.501 | Overall |  |  | 1.1060 | 1.4910 | 1.3676 | 1.6068 |
|  |  |  |  |  |  | undeg | 2.0885 | 3.6201 | 1.6861 | 2.2880 | 1.9465 | 3.2636 |
|  |  |  |  |  |  | tempdeg | 1.8115 | 2.9253 | 1.5634 | 2.2850 | 1.4055 | 3.4480 |
|  |  |  |  |  |  | specdeg | 0.0355 | 1.3142 | 0.0686 | 2.7332 | 0.7508 | 3.4193 |

